# The *Pseudomonas aeruginosa* Tse4 toxin assembles ion-selective and voltage-sensitive ion channels to couple membrane depolarisation with K^+^ efflux

**DOI:** 10.1101/2025.02.14.638235

**Authors:** Jessica Rojas-Palomino, Carmen Velázquez, Jon Altuna-Alvarez, Amaia González-Magaña, Maialen Zabala-Zearreta, Matthias Müller, María Queralt-Martín, Antonio Alcaraz, David Albesa-Jové

**Author notes:** To whom correspondence should be addressed: Antonio Alcaraz and David Albesa-Jové.

## Abstract

*Pseudomonas aeruginosa* employs the Type VI secretion system (T6SS) to outcompete other bacteria in its environment. Among the effectors secreted by the T6SS of *P. aeruginosa* PAO1, Tse4 is known for its potent antibacterial activity. This study elucidates the molecular function of Tse4, which promotes cell depolarisation in competing bacteria. Our results show that Tse4 spontaneously incorporates into lipid monolayers and forms multiionic channels in planar bilayers, with either ohmic conduction or diode-like rectifying currents and a preference for cations over anions. These observations allow us to propose a model of action whereby Tse4 channels couple cell depolarisation with K^+^ efflux. These insights into Tse4’s pore-forming activity enhance our understanding of bacterial competition and exemplify a finely tuned antibacterial strategy, coupling its ability to cause membrane depolarisation with potassium efflux that synergises with other T6SS effectors. These results highlight the sophistication of *Pseudomonas aeruginosa*’s competitive arsenal.

## Introduction

Bacterial competition is a fundamental aspect of microbial ecosystems, playing a pivotal role in shaping the dynamics of bacterial communities and influencing ecological and pathogenic processes. Understanding the mechanisms underlying this bacterial competition is of paramount importance, as it unveils the intricate strategies employed by microorganisms to gain a competitive advantage in their habitats. In this context, the present research focuses on elucidating the molecular function of Tse4, a <u>T</u>ype VI <u>s</u>ecretion <u>e</u>xported effector produced by *Pseudomonas aeruginosa*, a versatile opportunistic pathogen known for its ability to adapt, survive, and persist in diverse environments.^1^

The Type VI secretion system (T6SS) is a complex molecular machine, structurally and evolutionary related to the bacteriophage tail and spike, that many gram-negative bacteria use to inject effector proteins directly into neighbouring bacterial or eukaryotic cells.^2–8^ The *P. aeruginosa* genome contains three independent T6SS clusters (H1, H2, and H3-T6SS).^9,10^ *P. aeruginosa* PAO1 deploys the H1-T6SS in response to an interspecies^11^ or intraspecies^12^ attack, a crucial role in bacterial competition.

The effectiveness of the *P. aeruginosa* H1-T6SS in bacterial competition lies in the multitude and variety of the antiprokaryotic toxins it injects into competing bacteria. Therefore, unravelling the molecular functions of these toxins is crucial for gaining insights into this bacterial competition mechanism. The H1-T6SS of *P. aeruginosa* PAO1 is known to deliver eight antiprokaryotic toxins, targeting various cellular components and functions. These include (i) the cell wall peptidoglycan, with Tse1 exhibiting peptidase activity,^13,14^ and Tse3 displaying muramidase activity;^15,16^ (ii) NAD(P)+, impacted by Tse6 possessing glycosidase activity;^17^ (iii) the DNA, influenced by Tse7 with DNase activity;^18^ (iv) protein biosynthesis, modulated by Tse8;^19,20^ and (v) the membrane, affected by Tse4^6,21^ and Tse5.^22,23^

In the present study, we investigate the molecular function of Tse4 (PA2774), one of the most potent antiprokaryotic H1-T6SS effectors of *P. aeruginosa* PAO1 known to date.^24^ Tse4 associates with <u>h</u>aemolysin <u>c</u>o-regulated <u>p</u>rotein 1 (Hcp1) for H1-T6SS-dependent delivery into target cells.^6,25^ Along with its cognate immunity protein Tsi4, Tse4 can also be found in the inner membrane of *P. aeruginosa.*^26^ Previous in vivo studies showed that Tse4 induces relatively sophisticated membrane permeabilisation regarding ion specificity and size exclusion.^21^ Thus, it was shown that intoxication by Tse4 promoted cell growth inhibition that critically depended on electrolyte composition, being stronger in sodium and lithium chloride than in equimolar solutions of potassium chloride or sucrose. Also, several small solutes between 300 – 700 Da (0.45 – 0.65 nm equivalent hydrodynamic radius) could not access the cytoplasm of Tse4-intoxicated cells. Taking all these results into account, it was hypothesised that Tse4 promotes the formation of relatively narrow ion-selective membrane pores.^21^ However, this hypothesis was not verified at the molecular level, probably due to the difficulty in expressing and purifying Tse4 for in-vitro studies.^25^ Here, we successfully produced Tse4 by exploiting a novel purification system based on the encapsulation properties of Tse5^23^, demonstrating its potential biotechnological application for expressing toxic and/or hydrophobic proteins and allowing us to perform the first-ever biophysical study of the Tse4 function at the molecular level.

Our data directly demonstrate that Tse4 is a pore-forming toxin that integrates spontaneously into lipid monolayers, forming ohmic or rectifying channels in planar bilayers. Detailed electrophysiological analyses also reveal that Tse4-induced pores exhibit a general preference for cations that can range from mild to substantial selectivity, including a certain degree of chemical specificity. The fact that Tse4-poration activity does not involve detergent-like mechanisms disintegrating the membrane integrity aligns with in vivo data showing that Tse4-induced toxicity causes cell depolarisation and bacteriostasis.^21^ Remarkably, the finding of non-ohmic Tse4-induced pores that open preferentially at positive potentials suggests that they may operate as outward rectifying channels, potentially playing a role in the modulation of action potentials, as observed with other toxins known to influence membrane excitability.^27–29^

## RESULTS

### Heterologous expression of a Tse5^ΔCT^ -Tse4 chimera

To overcome the challenges of producing Tse4 due to its hydrophobicity and toxicity, we explored the encapsulating capacity of Tse5, a *P. aeruginosa* H1-T6SS-dependent Rearrangement hot spot (Rhs) toxin.^6,30^ Over a decade of research into bacterial Rhs toxins has revealed tremendous insight into their structures and mechanisms of action. Examples of bacterial Rhs toxins involved in the intercellular competition include insecticidal toxin complexes (Tc),^31,32^ Gram-negative T6SS-associated Rhs toxins,^33–35^ and the Gram-positive wall-associated protein A (WapA).^36^

Rhs polymorphic toxins contain tyrosine/aspartate repeats (YD-repeats)^37^ that assemble into a β-sheet that spirals to form a barrel-like/cocoon structure.^33,34,38–40^ A recent cryo-EM structure revealed that Tse5 is organised in three fragments.^23^ The N-terminal fragment contains a <u>p</u>roline-<u>a</u>lanine-<u>a</u>lanine-<u>p</u>roline (PAAR)−like motif that presumably targets the effector to the T6SS. This fragment results from auto-cleaved between Lys47 and Pro48 residues by a yet-to-be-discovered mechanism. The central fragment assembles this cocoon (Tse5-Shell), encapsulating the C-terminal toxic fragment (Tse5-CT). Cleavage of the toxic fragment is vital for toxin activation. This cleavage is mediated by a conserved aspartyl protease domain characterised by a DPxGx_19_DPxG motif and located at the C-terminal end of the Tse5-Shell fragment.^23^ The toxicity of Tse5-CT is attributed to the assembly in the cytoplasmic membrane of competing bacteria of ion-selective proteolipidic pores. This ion channel activity of Tse5-CT causes cell depolarisation and, ultimately, bacterial death.^22^

Based on Tse5’s capacity to encapsulate the pore-forming toxin Tse5-CT, we decided to investigate whether Tse5-Shell could be used to encapsulate Tse4, which contains several predicted transmembrane regions (**Supplementary Fig. 1**). The Tse5-Shell’s cavity volume is ∼32,000 Å^3^, which should be sufficient to encapsulate the 19.2 kDa Tse4 protein, which has a calculated dry volume of 23,491 Å^3^.

To this end, we have engineered a chimeric protein based on Tse5, where the Tse4 sequence replaces its Tse5-CT fragment (**Fig. 1A, Supplementary Fig. 2**). Tse4 is inserted after Leu1168 residue of Tse5, which is where the aspartyl protease domain of the Tse5-Shell cleaves. The chimeric Tse5^ΔCT^-Tse4 protein contains a poly-His tag at the N-terminus for protein purification by Ni-NTA affinity chromatography, and it also contains mutations K47G and P48A that inhibit auto-cleavage of its N-terminal fragment. Remarkably, <u>s</u>ize <u>e</u>xclusion <u>c</u>hromatography-<u>s</u>mall <u>a</u>ngle <u>X</u>-ray <u>s</u>cattering (SEC-SAXS) indicates Tse5-CT deletion mutant (Tse5^ΔCT^) and Tse5^ΔCT^-Tse4 have comparable hydrodynamic behaviour, as indicated by having comparable SEC elution profiles and radius of gyration (*R*_g_ = 36.5 Å and 36.2 Å for Tse5^ΔCT^ and Tse5^ΔCT^-Tse4, respectively, as calculated from the Pair distance distribution function [*P*(*r*)]; **Table 1**, **Fig. 1B)**. Furthermore, when running Tse5^ΔCT^-Tse4 on a denaturing <u>s</u>odium <u>d</u>odecyl <u>s</u>ulphate-<u>p</u>oly<u>a</u>crylamide <u>g</u>el <u>e</u>lectrophoresis (SDS-PAGE), Tse4 appears as a fragment around the ∼20 kDa protein marker, indicating the Tse5^ΔCT^ aspartyl protease domain remains active (**Fig. 1C**). Following Tse5^ΔCT^-Tse4 expression and purification, we can separate Tse4 from Tse5^ΔCT^ by denaturation with 6 M urea and precipitation with ammonium phosphate (**Fig. 1C**, see Methods section for experimental details). Altogether, this data indicates that Tse4 is cleaved and remains encapsulated inside Tse5^ΔCT^.

**Figure 1.**
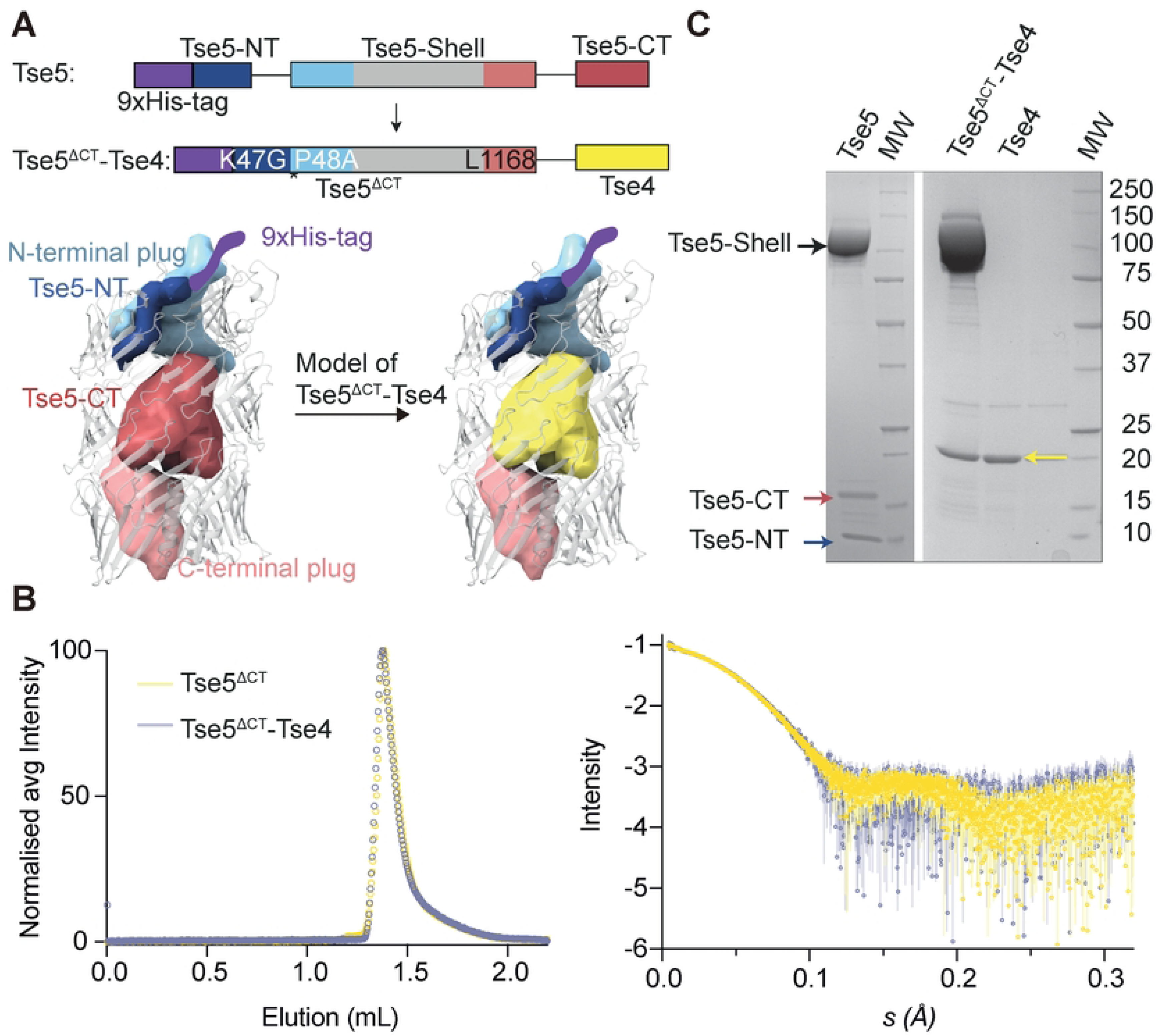
Heterologous expression of a Tse5^ΔCT^-Tse4 chimera. **A.** The top panel shows a schematic representation of Tse5 and Tse5^ΔCT^ -Tse4 constructs. Tse4 replaces the Tse5-CT fragment. The Tse5^ΔCT^-Tse4 construct contains a double point mutation (K47G-P48A) that inhibits cleavage between residues G47 and A48. The bottom panel shows a predicted 3D structure of Tse5 and Tse5^ΔCT^ -Tse4 constructs. The construct is designed so that Tse4 is encapsulated inside the Tse5 shell/cocoon structure. **B.** SEC-SAXS analysis of Tse5 and Tse5^ΔCT^-Tse4 proteins. The left panel shows the normalised SEC signal profile for both proteins, indicating they have very similar elution properties. The right panel plot represents the SAXS data for the two proteins. **C.** SDS-PAGE of purified wild-type Tse5 (left) showing that WT Tse5 auto-cleaves, resulting in three fragments, Tse5-Shell, Tse5-CT and Tse5-NT. On the right is the SDS-PAGE of Tse5^ΔCT^-Tse4 and Tse4 following its separation from the Tse5^ΔCT^ fragment.

**Table 1.**
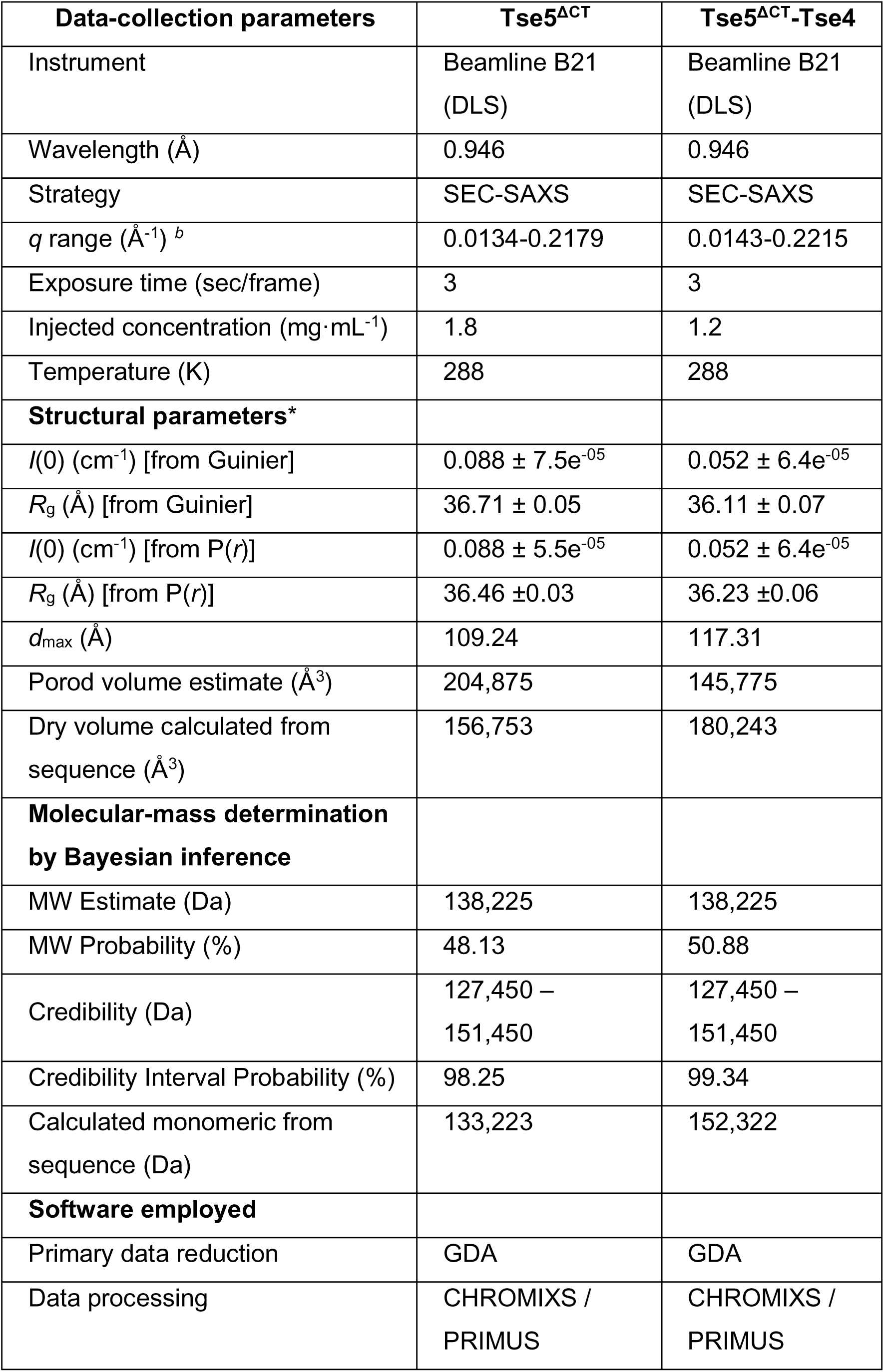
Summary of SEC-SAXS data analysis for Tse5^ΔCT^ and Tse5^ΔCT^-Tse4.

### Tse4 inserts spontaneously into model membranes to assemble narrow membrane pores

Previous cellular studies demonstrated that Tse4 leads to impaired growth in *P. aeruginosa*^21^, displaying certain sensitivity to specific monovalent cations, particularly Na^+^ and Li^+^ ions. Also, these results provided evidence that Tse4 disrupts the proton motive force by affecting the membrane potential but not pH homeostasis, allowing the passage of ions while excluding larger molecules.

Although the above findings suggested that Tse4 functions by facilitating the formation of ion-selective membrane pores, there was no direct evidence to exclude alternative potential pathways involving other cellular components. To address these questions and provide insight into its molecular mechanism, we first evaluated the capacity of Tse4 to spontaneously partition into the hydrophobic core of lipid monolayers assembled from an *E. coli* polar lipid extract using the Langmuir-Blodgett balance^41^ (refer to the Methods section for experimental details). This method detects the insertion of a protein into the hydrophobic core of a monolayer by monitoring the change in lateral pressure (Δ*Π*) from an initial lateral pressure (*Π*_0_). As the initial lateral pressure increases, the rate of protein insertion decreases until it reaches a critical lateral pressure (*Π*_c_) where protein is not able to insert and therefore Δ*Π* = 0. In the biological membrane’s outer monolayer, the lipid packing typically generates lateral surface pressures between 30 to 35 mN/m.^42,43^ Therefore, when the critical lateral pressure falls within this range, it indicates that the protein is spontaneously integrating into the hydrophobic core of the lipid monolayer.

The incorporation of Tse4 into lipid monolayers with diverse *Π*_0_ resulted in a critical lateral pressure of 31.1 mN/m, therefore indicating Tse4 spontaneously integrates into the hydrophobic core of the lipid monolayer (**Fig. 2A, B**). Remarkably, Tse5^ΔCT^-Tse4 and Tse5^ΔCT^ display similar *Π*_c_ that are below the 30 mN/m threshold, indicating that neither of them spontaneously integrates into the hydrophobic core of the lipid monolayer (*Π*_c_ values for Tse5^ΔCT^-Tse4 and Tse5^ΔCT^ are 26.51 and 27.17, respectively).

**Figure 2.**
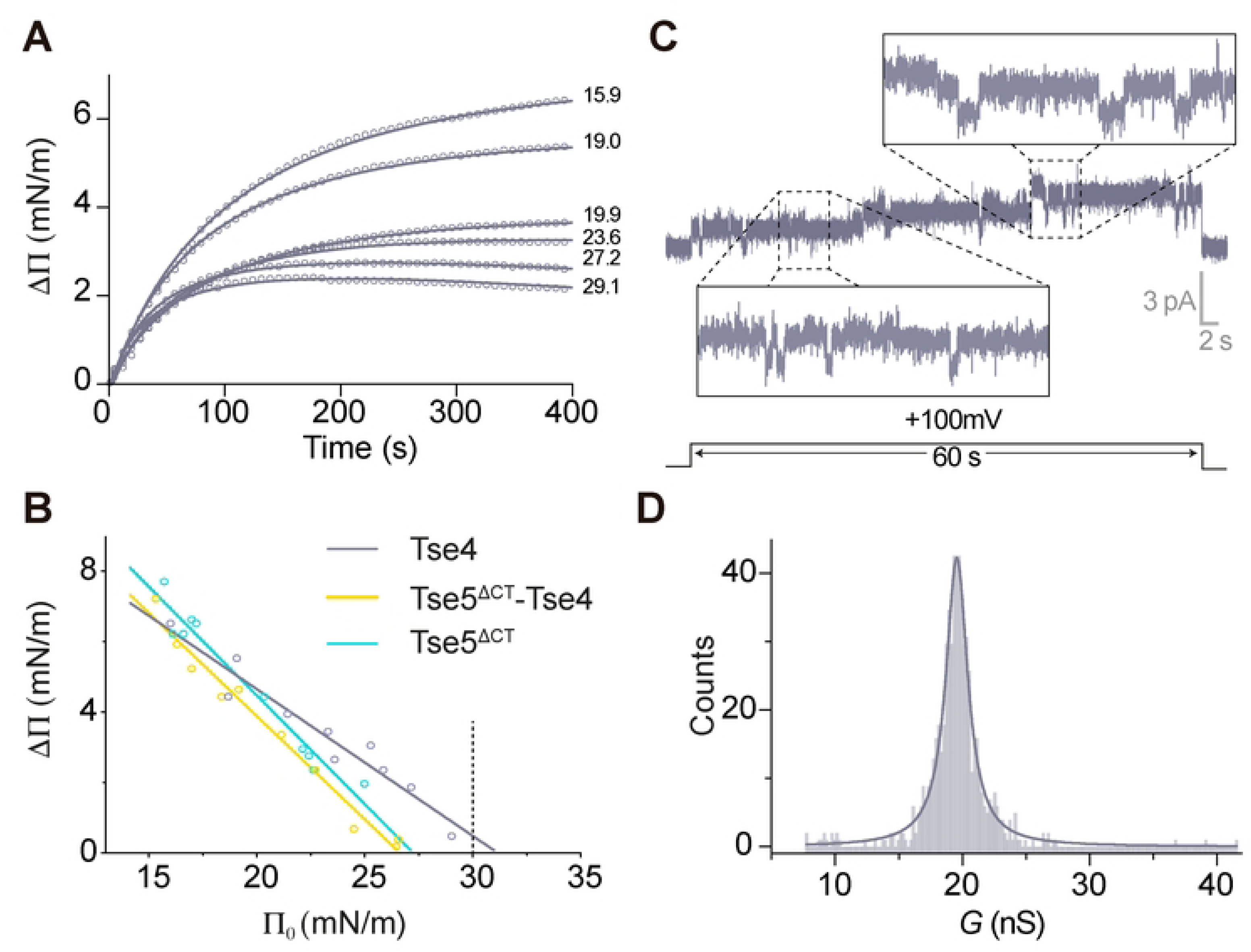
Tse4 inserts spontaneously into model membranes to assemble narrow membrane pores. **A.** Representative Langmuir-Blodgett balance data showing the lateral pressure increase on lipid monolayers after the addition of Tse4 at time 0. Initial lateral pressures (*Π*_0_) in mN m^−1^ for representative experiments are indicated above each curve. **B.** Plot of lateral pressure increases (Δ*Π*) as a function of initial lateral pressure (*Π*_0_) for Tse4, Tse5^ΔCT^-Tse4, and Tse5^ΔCT^ (*n* = 10). A maximal insertion pressure (*Π*_c_) of 31.08 (Tse4), 26.53 (Tse5^ΔCT^-Tse4), and 27.17 mN m^−1^ for Tse5^ΔCT^ has been determined by extrapolating the fitted curve to Δ*Π* = 0. The dotted line indicates the threshold value of lateral pressure consistent with unstressed biological membranes. The equations obtained from the linear regression analysis are y = −0.4148*x* + 12.89 (*R*-squared = 0.94), *y* = −0.5832*x* + 15.47 (*R*^2^-squared = 0.98), *y* = −0.6141*x* + 16.68 (*R*-squared = 0.96) for Tse4, Tse5^ΔCT^-Tse4, and Tse5^ΔCT^, respectively. **C.** Representative current traces showing small current jumps recorded at a constant voltage of 100 mV in symmetrical 150 mM KCl. The raw current variance is overlaid with a 4th-order polynomial smoothing function. **D.** Frequency distribution analysis of 644 independent current jumps obtained at 100 mV was conducted using a bin width of 0.2 p*S*. The resulting dataset, comprising 170 data points, was fitted to a Lorentzian function (*R*² = 0.97), indicating a peak in the distribution at 19.6 ± 1 pS.

The capacity of Tse4 to integrate into a model membrane prompted us to investigate its ability to form membrane pores. To this end, we implement a modified solvent-free Montal-Mueller technique^44^ (see Methods section for experimental details). First, we assemble a lipid bilayer using the *E. coli* polar lipid extract and a concentration of 150 mM KCl at pH 7.4 on both chambers. Following the addition of 0.4 µM of Tse4 in the *cis* chamber, we observe ion-channel-like activity in the form of relatively stable currents but also exhibiting transitions between different conductive levels with a diversity of lifetimes (**Fig. 2C**). Since from currents recordings we cannot discriminate between the collective action of clusters of small units and potential individual wide pores we considered traces where the current was variable and calculated the conductance increments (ΔG = ΔI/V) associated to each current jump ΔI. As expected from the random nature of current jumps, we obtain a Gaussian distribution with a peak located at G ∼ 20 ± 3 pS (**Fig. 2D**), which is comparable to values measured under similar conditions (salt concentration and membrane composition) for channels of known structure such as Gramicidin A (gA) (G ∼ 30 pS, r ∼ 0.4 nm)^45^ and the lowest level of the antibiotic peptide Alamethicin, namely L0 (G ∼ 50 pS, r ∼ 0.7 nm)^46^. Hence, an estimation of r ∼ 0.5 nm for Tse4-induced pores seems reasonable and agrees satisfactorily with previous studies showing that molecules larger than 300 Da, about 0.5 nm of hydrodynamic radius, were unable to access the cytoplasm of Tse4-intoxicated cells^21^.

### Tse4 forms a variety of cation-selective protein pores with mild cation specificity

Next, we measured the applied voltage needed to cancel the current (the so-called <u>R</u>eversal <u>P</u>otential or RP) in experiments with Tse4 under a concentration gradient of 250/50 mM of different salts. In all studied salts (KCl, NaCl, LiCl), we see a variety of cation-selective channels (RP < 0 always), as shown in the histograms of **Fig 3A**. The higher dispersion is found in KCl, where the maximum probability is around RP = −20 mV. Interestingly, the peaks for NaCl and LiCl appear at RP = −22 mV and −13 mV, respectively. Considering that the theoretical limit (Nernst potential setting the ideal maximum selectivity) for 250/50 mM gradient is RP ∼ −41mV, our data suggest Tse4-induced membrane pores have a multi-ionic character but with a marked preference for cations. Indeed, this image appears when RP measurements are turned into ion permeability ratios (P_+_/P_-_) using the Goldman-Hodgkin-Katz flux equation (GHK). Tse4-induced pores in model *E. coli* cytoplasmic membrane showed a P_Na_+/P_Cl^-^_ = 4.4 ± 1.6, P_K_+/P_Cl^-^_ = 4.0 ± 1.9 and P_Li_+/P_Cl^-^_ = 2.4 ± 0.9 (**Fig. 3B**). Notice that P_K_+/P_Cl^-^_ = 4.0 ± 1.9 correspond to ohmic pores. As shown in the next section, Tse4 in KCl also induces the formation of non-ohmic channels with a higher selectivity (P_K_+/P_Cl^-^_ ∼ 19) (indicated as ‘KCl rectifying’ in **Fig. 3A-B**).

**Figure 3.**
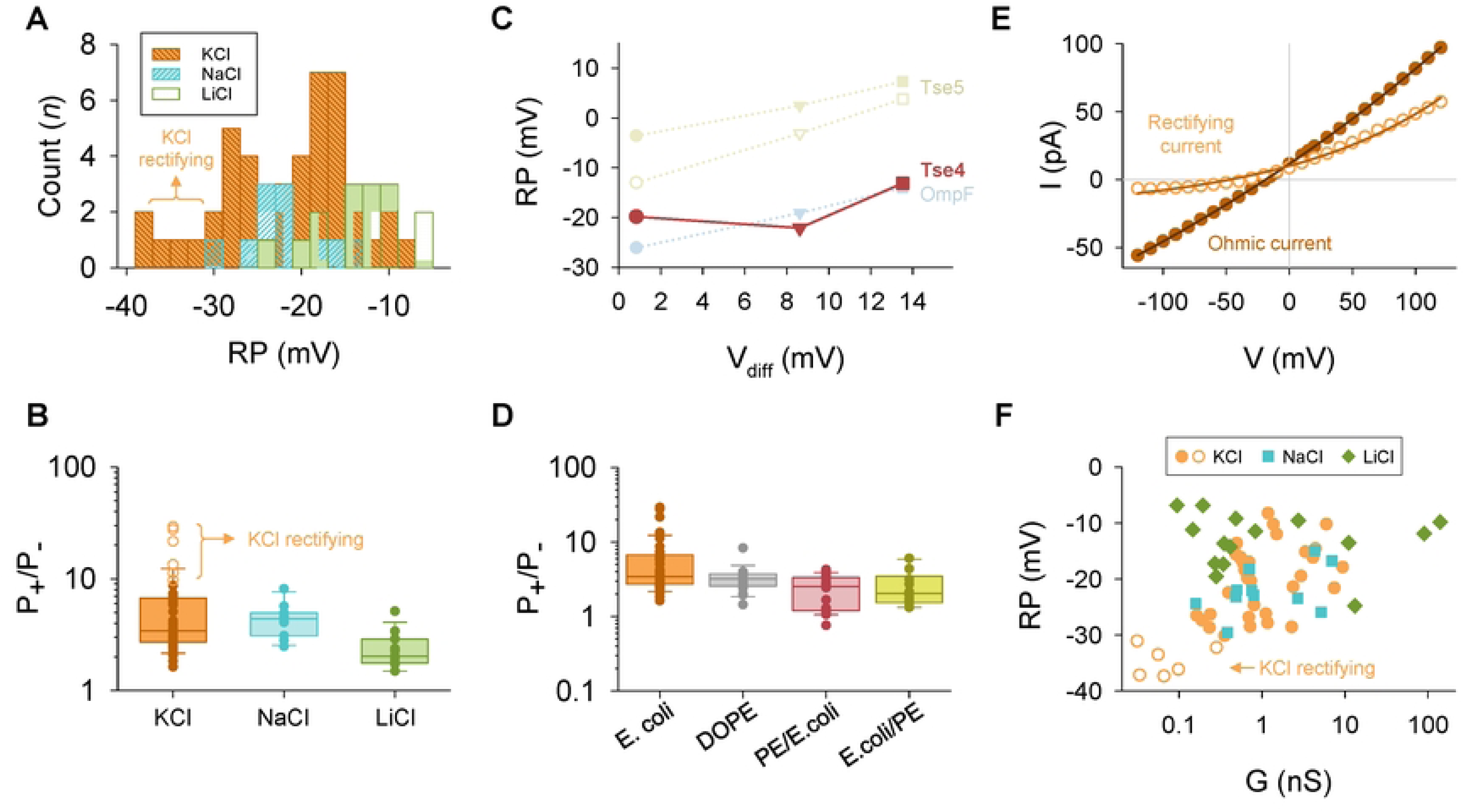
Tse4 forms a variety of cation-selective pores with mild cation specificity and diverse voltage sensitivity. **A.** Histograms illustrating the reversal potentials (RPs) for Tse4-induced channels in KCl (orange, n = 43), NaCl (blue, n = 11) or LiCl (green, n = 15). The frequency distribution analysis was performed with a bin width of 2 mV. Rectifying pores in KCl display the most negative RPs. **B.** Box and whiskers plot displaying the P_+_/P_-_ ratios in KCl, NaCl and LiCl derived from the experiments presented in panel A. Here and elsewhere, the boundary of the box closest to zero indicates the 25^th^ percentile, a line within the box marks the median, and the boundary of the box farthest from zero indicates the 75^th^ percentile. Whiskers (error bars) above and below the box indicate the 90^th^ and 10^th^ percentiles. Rectifying pores in KCl exhibit the maximum P_+_/P_-_ values. **C.** RP measured for Tse4 in E. coli and compared with Tse5 and OmpF in KCl (circles), NaCl (triangles) or LiCl (squares) plotted versus the corresponding diffusion potential (V_diff_) for each salt. For Tse5, results are shown for charged (solid symbols) and neutral (open symbols) lipids. RP for Tse4 in KCl only includes ohmic channels, as there are no rectifying currents in other salts. **D.** Box and whiskers plot displaying the P_+_/P_-_ ratios measured in Tse4 in KCl using different lipid compositions, as indicated. **E.** Example current-voltage curves depicting the electrical behaviour of ohmic (filled circles) and rectifying (open circles) pores induced by Tse4 in KCl and E. coli lipid mixture. The solid lines represent a fit with an empirical equation used to categorize I-V curves as ohmic or rectifying (see Methods for details). **F.** Scatter plot of the RP as a function of the conductance measured in KCl (filled circles correspond to ohmic currents while open circles indicate rectifying currents), NaCl, and LiCl, derived from the experiments presented in panels A and B. In all panels, experiments were performed in a 250 / 50 mM salt gradient.

Concerning a possible channel specificity among cations, note that the ionic selectivity is a property of the system that depends on the channel and the electrolyte solution flowing through it. Consequently, the measured RP includes the diffusion potential (DP) arising from the intrinsic differences in ionic diffusivities between cations and anions^47^. For a 250/50 mM gradient, DP can be calculated via Planck’s equation using tabulated free solution values of ionic diffusion coefficients^47^ yielding DP ∼ 0.8 mV for KCl (note that K^+^ and Cl^-^ ions have very similar diffusion coefficients), DP ∼ 8.6 mV for NaCl, and 13.5 mV for LiCl (note that for the considered gradient all DP values are positive whereas all measured values are negative). If the channel selectivity were non-specific, RP measured for salts of different cations should differ approximately in the difference between their respective DP, as it is the case in the channels formed by Tse5^48^ or from the bacterial porin OmpF of *Escherichia Coli^49^*. These channels are considered non-specific and yield an approximate straight line when plotting their RP vs. DP (**Fig. 3C**). However, this is not the case in Tse4-induced channels: the difference in RP in KCl vs NaCl should be +8 mV, and it is –2 mV. Likewise, the difference in RP in KCl vs LiCl should be +13 mV, but it is only +7 mV, so the plot RP vs. DP does not show a straight line (**Fig. 3C)**. Considering this, we conclude that Tse4-induced pores have a greater channel discrimination for Na^+^ and Li^+^ than for K^+^.

Tse4 channel selectivity may be mainly controlled by the protein characteristics or could otherwise be influenced by the membrane lipid charge, as is the case of a variety of pore-forming proteins such as viroporins,^50^ cell-penetrating peptides^51–53^ or toxins including Tse5.^22,48,54,55^ To test this possibility, we measured the RP of Tse4-induced channels in a neutral lipid, DOPE, and in asymmetric membranes formed of *E. coli* mixture on one side and DOPE on the other side of the membrane. Interestingly, we observe no relevant effect of lipid charge in any of the membrane variations tested, obtaining cation selective channels similar to that in *E. coli* even in the fully neutral lipid (P_K_+/P_Cl^-^_ = 3.3 ± 1.5) (**Fig. 3D**). Based on this evidence, we presume that Tse4-induced pores are basically protein channels where lipid molecules are not structurally involved in the channel architecture.

### Non-Ohmic behaviour and rectification in Tse4 channels and its implications for selectivity and conductance

Selectivity experiments contain more information than the RP corresponding to I = 0. Thus, we can also consider the full current-voltage (I-V) relationship, where we observed mainly linear I-V curves. However, for KCl in *E. coli* lipid mixtures we detected some that seemed to deviate from linearity, so we formally classified the recorded currents into ohmic (resistor-like conduction) and non-ohmic (asymmetric diode-like conduction) based on an ideality factor (see Methods section for details) (**Fig. 3E and Supplementary Fig. 3**). Our data shows that, out of 44 independent channels, 86% are ohmic (n = 38) and 14% are rectifying (n = 6) (**Supplementary Fig. 3**). Interestingly, all non-ohmic pores act as outward rectifying channels, given that they open at positive potentials allowing mostly cation efflux^56^. Rectifying I-V curves are usually associated with structural inhomogeneities in the system, either in terms of pore geometry (i.e., conical pores) or charge distribution^57^.

In all cases, the measured RP can be associated with the experimental G obtained as the I/V slope if the curve is ohmic or as the local I/V quotient in the I = 0 region for rectifying I-V curves. For all the electrolytes under investigation, we observe a considerable dispersion in the RP vs G representation (**Fig. 3F**). For KCl and NaCl, it is tempting to hint a relationship between the greater selectivity and the associated value of G, suggesting that channel enhanced selectivity (higher values of RP) is attained by narrowing the permeation pathway for ions (lower values of G). However, this is not the case for LiCl, where smaller values of G correspond to lower values of RP, indicating that the proximity of protein charges to the pore eyelet is not the only factor ruling ion selectivity, but other mechanisms exist, such as some specificity derived from residue-ion interactions^58,59^.

The case of KCl is particularly interesting because the high dispersion found in RP values can be rationalized in terms of ohmic and rectifying channels. Thus, while low and moderate RP values correspond to weakly selective (P_K_+/P_Cl^-^_ ≤ 10) ohmic channels (orange filled circles in **Fig. 3F and Fig. 3B**), slightly higher RP (P_K_+/P_Cl^-^_ ∼10-30) values are linked to rectifying channels (open orange circles in **Fig. 3F and Fig. 3B**).

### A conceptual model for Tse4-induced changes in cell membrane potential

Potassium is the most abundant intracellular cation in all living organisms, where it is required for numerous basic cellular functions, including regulating intracellular pH, governing the magnitude of the transmembrane electrical potential, and balancing turgor/osmotic pressure.^60^ Therefore, the ability of bacteria to regulate intracellular potassium (K⁺) levels is critical for maintaining homeostasis, surviving environmental stresses, and adapting to hostile host environments. Bacteria employ multiple potassium transport systems, each tailored to specific environmental conditions, to achieve this regulation. These systems not only play fundamental physiological roles but also contribute to pathogenesis and antimicrobial resistance.^61^

Previous cellular studies showed that Tse4 induces a potassium flux from the cytoplasm to the extracellular milieu.^21^ In principle, such externalisation of positive charge should lead to a hyperpolarised state in which the charge imbalance would dramatically alter the intracellular pH. However, the actual situation was quite different. Membrane depolarisation was proven using potential-sensitive fluorescent dyes, and pH-sensitive fluorescent proteins showed no changes in pH in Tse4-intoxicated cells.^21^ Given that, we postulate that a more subtle mechanism could operate in which Tse4-induced pores disturb cellular homeostasis by playing a dual role in terms of cation specificity and voltage-dependent conduction as shown in **Fig. 4**.

**Figure 4.**
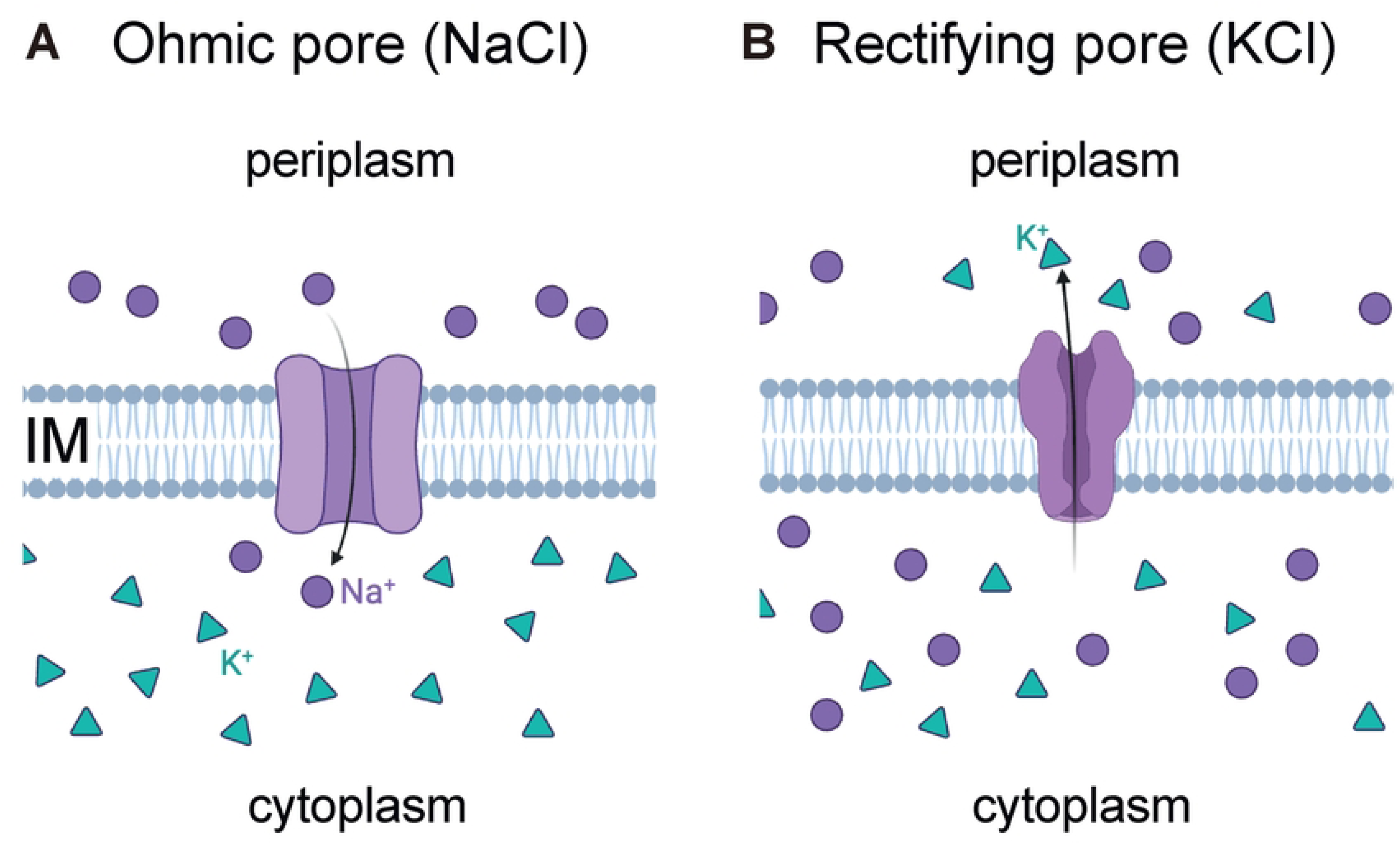
A conceptual model for Tse4-induced changes in cell membrane potential. Tse4 forms ion-selective pores in bacterial membranes, exhibiting mild specificity for Na⁺ and Li⁺ over K⁺. These pores facilitate Na⁺ influx (**4A**) and K⁺ efflux (**4B**), driven by concentration gradients and membrane potential. Tse4-induced K⁺ efflux is enhanced under depolarised conditions due to rectifying pore conduction, maintaining cytoplasmic electroneutrality without affecting pH (Fig. 4B). This model integrates observations of K⁺ efflux, membrane depolarisation, and unchanged pH, highlighting Tse4’s role in disrupting membrane potential and sensitising bacteria to other T6SS effectors.

Our electrophysiological studies demonstrate that Tse4 forms ion-selective pores in bacterial membranes, displaying a relative preference for monovalent cations over anions, including mild Na⁺ and Li⁺ specificity compared to K⁺. These findings correlate well with the previous in vivo work,^21^ which showed that Tse4 toxicity in *Pseudomonas aeruginosa* depends on the type of extracellular salt, particularly with increased sensitivity to sodium and lithium ions compared to potassium or sucrose. Considering this, we hypothesise that Tse4-induced cell depolarisation could be due to the influx of Na^+^ through the ohmic pores created in the membrane (**Fig. 4A**). To justify this premise, we consider not only the cationic selectivity observed in Tse4 experiments (including a slight preference for Na^+^ over K^+^) but also the asymmetric situation of these ionic species in the respective cellular compartments. At resting potential Na^+^ ions accumulate in the periplasm while being excluded from the cytoplasm due to the action of Na+/K+-ATPase. Accordingly, in the case of pore formation, both the concentration gradient and the resting potential push Na^+^ ions towards the cytoplasm (**Fig. 4A**), in contrast to K⁺ cations and Cl^-^ anions that are mostly unaffected because the membrane resting potential is relatively close to their respective equilibrium potentials. Remarkably, we have also observed that in the case of KCl, Tse4-induced pores may exhibit non-ohmic (rectifying) conduction. These rectifying pores are highly selective to cations (see **Fig. 3B**) so that they could facilitate K⁺ efflux while rejecting Cl^-^ influx, with the efflux rate increasing in correlation with less negative membrane potentials—consistent with in vivo observations.

With the available biophysical and in vivo data, we speculate that, when a few Tse4 molecules are delivered into the target bacterium, some of them form ohmic channels that allow instant Na^+^ influx while others create outward rectifying channels that are opened when the membrane becomes depolarised (**Fig. 4B**). Within this simplified conceptual model, the influx of Na^+^ is balanced in terms of charge by the efflux of K^+^ so that no change in proton concentration or chloride influx is required to maintain electroneutrality in the cytoplasm. As a result, all experimental evidence (K^+^ efflux, membrane depolarisation, and no pH change) could be included in a single unified mode of action.

It should be noted that such a conceptual model may be oversimplified because it focuses only in cation transport disregarding the possibility of direct interactions between Tse4 and other cellular components, such as the endogenous bacterial Na^+^ and K^+^ channels^62,63^ or the voltage dependence of the Na+/K+-ATPase.^64^ Furthermore, Tse4 synergises with other effectors to enhance antibacterial activity. Tse4’s disruption of the membrane potential and K^+^ efflux could also explain the observed increased sensitivity of target cells to the activity of other T6SS effectors, such as Tse1 and Tse3, which degrade peptidoglycan, and Tse6, a NAD+ glycohydrolase. The loss of the membrane potential and K^+^ efflux could activate <u>p</u>roton <u>m</u>otive <u>f</u>orce (PMF)-sensitive autolysins^65^, accelerating the structural degradation caused by cell wall-targeting effectors, while concurrently inhibiting PMF-dependent transporters, exacerbating the metabolic stress induced by Tse6.^21^

## CONCLUSION

In this study, we successfully expressed and purified the Tse5ΔCT-Tse4 chimera, overcoming the inherent challenges posed by the hydrophobicity and toxicity of Tse4. This was achieved using the encapsulating properties of Tse5, which allowed us to isolate Tse4 for further biophysical and electrophysiological studies. By employing this chimeric expression system, we have investigated the pore-forming activity of Tse4 in vitro and its role in bacterial competition. Our results show that Tse4 spontaneously incorporates into lipid monolayers and forms multiionic channels in planar bilayers, showing either ohmic conduction or diode-like rectifying currents with mild to significant cation selectivity. Based on this, we hypothesise that Tse4-induced pores disturb cellular homeostasis by playing a dual role in terms of cation specificity and voltage-dependent conduction.

## METHODS

### Construct design, protein expression and purification

The gene coding for Tse4 (PA2774) was synthesised by GenScript (GenScript, NJ, USA) and cloned into a pET29a(+) vector (pet29a(+)::K47G-P48A) that derives from the parental vector pet29a(+)::9xhis-Tse5. The last plasmid includes, between the NdeI and HindIII restriction sites, a 5’ 9xHis-tag and a tobacco etch virus protease cleavage site (ATGGGCAGCAGCCATCATCATCATCATCATCATCATCACAGCAGCGGCGAAAACCTGTATTTTCAGGGCGGATCC), followed by the coding sequence of Tse5 (PA2684). To avoid N-terminal cleavage of Tse5, a GenScript derived mutation led to the new parental vector pet29a(+)::K47G-P48A, containing two single point mutations of residues K47 and P48 to glycine and alanine, respectively. The protein sequence of Tse4 was cloned into pet29a(+)::K47G-P48A exchanging Tse5-CT toxin sequence. The final pet29a(+)::*tse5*^ΔCT^-*tse4* plasmid codes for the construct Tse5^ΔCT^-Tse4 (Supplementary Table 1 and Supplementary Data 1).

*Escherichia coli* Lemo21(DE3) cells were transformed with pET29a(+)::*tse5*^ΔCT^*-tse4* plasmid and grown overnight at 37 °C in shaking conditions within flasks containing 200 mL of LB media supplemented with 50 μg/mL kanamycin, 34 μg/mL chloramphenicol and 2 mM rhamnose. For Tse5^ΔCT^-Tse4 overexpression, bacterial cultures were diluted to OD_600_ value of 0.1 with 2 L of fresh LB medium supplemented with both antibiotics at the same concentrations, but lacking rhamnose. When cells reached an OD_600_ value of 0.7 after growing at 37 °C in shaking conditions, protein expression was induced by adding isopropyl β-D-1-thiogalactopyranoside (IPTG) at the final concentration of 1 mM. Bacterial cultures were then left overnight with agitation at 18 °C. Finally, cells were pelleted and stored at −80 °C for later use.

Pellet from 2 L of bacterial culture was resuspended in 30 mL of 50 mM Tris–HCl pH 8.0, 500 mM NaCl, 20 mM imidazole (solution A) with 4 μL of benzonase endonuclease (Millipore, Sigma) and a tablet of protease inhibitor cocktail (cOmplete, EDTA-free, Roche). Cells were then disrupted by sonication for 3 minutes (continual cycles of 10 s ON and 59 s OFF with 60 % amplitude), and the suspension was ultra-centrifuged for 60 min at 125748 x g. The supernatant was filtered using a 0.2 μm syringe filter and then loaded into a HisTrap HP column of 5 mL (GE Healthcare) equilibrated with solution A to perform an immobilised metal affinity chromatography on a fast protein liquid chromatography system (ÄKTA FPLC; GE Healthcare). The column was washed with solution A at 0.3 mL/min until the value of absorbance at 280 nm was almost zero. Tse5^ΔCT^-Tse4 was eluted with 100% of 50 mM Tris–HCl pH 8, 500 mM NaCl and 500 mM imidazole (solution B) at 2 mL/min. Peak fractions were pooled, and protein was injected into a HiLoad Superdex 200 26/600 pg, previously equilibrated with 20 mM Tris–HCl pH 8, 150 mM NaCl and 2 mM DTT. Tse5^ΔCT^-Tse4 eluted as a single monodispersed protein but SDS–PAGE showed two major protein fragments, Tse5^ΔCT^ and Tse4 (**Fig. 1B, C**). The identity of the Tse4 band was confirmed by mass spectrometry (Supplementary Data 2). Fractions containing Tse5^ΔCT^-Tse4 (checked by SDS-PAGE) were pooled, and concentration was assessed by measuring absorbance at 280 nm (*ca.* yield: 30 mg/L).

Using Amicon centrifugal filter units of 30 kDa molecular mass cut-off (Millipore), Tse5^ΔCT^-Tse4 was concentrated up to 17 mg mL−1. To separate the two fragments, protein denaturation was achieved by diluting the concentrated protein in solution A with 8 M urea to a final concentration of 6 M. Once the protein was incubated for 30’ under constant shaking, it was loaded with a peristaltic pump (Cytiva) into a HisTrap HP column of 5 mL (GE Healthcare), previously washed and equilibrated with solution A containing 6 M urea. The column was washed with 20 mL of solution A at 0.3 mL/min, and unbound fractions (flowthrough) containing Tse4 were collected. Elution was then performed with 10 mL of solution B at 2 mL/min to recover the Tse5^ΔCT^ fragment. Flowthrough containing only Tse4 was precipitated with 1.5 M ammonium sulphate and 15 min centrifugation at 15000 × g. Precipitated Tse4 was washed twice with MiliQ, flash frozen with liquid nitrogen, lyophilised and then stored at −80 °C until use. The purity of the protein was confirmed by SDS–PAGE (**Fig. 1C**).

### Study the insertion of Tse5^ΔCT^-Tse4 and Tse4 in lipid monolayers using the Langmuir–Blodgett balance technique

The value of the critical lateral pressure (*Π*_c_) of Tse5^ΔCT^-Tse4 and Tse4 was measured by the Langmuir–Blodgett balance technique with a DeltaPi-4 Kibron tensiometer (Helsinki, Finland) to check whether an insertion into a lipid monolayer happens spontaneously. The temperature in each experiment was set at 25 °C using a water bath (JULABO F12). *E. coli* polar lipid extract (Avanti Polar lipids) dissolved in chloroform at 1 mg/mL was extended with a Hamilton microsyringe over 1.25 mL of the aqueous surface (5 mM Hepes pH 7.4, 150 mM NaCl) previously added into each circular trough (Kibron μTrough S system, Helsinki, Finland) of 2 cm in diameter. Starting from different initial surface pressure (*Π*_0_) of the lipid monolayer, within the interval of 15 to 30 mN/m, changes in surface pressure (ΔΠ) were recorded when the protein was injected into the aqueous subphase at a final concentration of 0.4 μM. Tse5^ΔCT^-Tse4 was dissolved in 20 mM Tris-HCl pH 8.0, 150 mM NaCl, 2 mM DTT, injecting the buffer alone as a control. Tse4 was dissolved in dimethyl sulfoxide (DMSO), so the control was performed by injecting DMSO alone. When the different ΔΠ were plotted as a function of Π_0_, data was fitted into a linear regression model to obtain the *Π*_c_ by extrapolation (y value when x = 0)

### Electrophysical study of the pore-forming activity for Tse4, Tse5 and OmpF in planar lipid bilayers

Planar lipid bilayers were formed by using a solvent-free modified Montal-Mueller technique^66^. In summary, bilayers were formed by apposition of lipid monolayers on a ∼150-μm-diameter hole within a 15-μm-thick Teflon film, separating two symmetrical compartments of a Teflon chamber: *cis* and *trans* (1.8 mL of internal volume each). 3% solution of hexadecane in pentane was used for hole pre-treatment. Membranes were formed from a natural polar lipid extract from *E. coli*, dioleoyl-phosphatidylethanolamine (DOPE) or a combination of both for Tse4 experiments, as indicated in each figure legend. In experiments with Tse5 and OmpF shown in **Fig. 3C**, lipids used were pure DOPE (Tse5, neutral lipid), a combination of dioleoyl-phosphoglycerol (DOPG) and DOPE at a ratio DOPE/DOPG 70:30 w/w (Tse5, charged lipid) or diphytanoyl phosphatidylcholine (OmpF, neutral lipid). All lipids were purchased from Avanti Polar Lipids (Supplementary Data 3). The chamber compartments were filled with KCl, NaCl or LiCl solutions at various concentrations indicated in each figure legend. All solutions were buffered by 5 mM Hepes at pH 7.4. After membrane formation and before protein addition, the absence of channel-like activity was verified by monitoring for 10 min with holding voltages ranging from -100 to +100 mV without detecting any non-zero current.

For experiments with Tse4, the protein dissolved in DMSO at a stock concentration of 2 mg/ml was added to the aqueous phase at one (*cis*) side of the bilayer to reach a final concentration of 0.4 µM. Tse5 and OmpF^67^ experiments shown in **Fig. 3C** were carried out as described previously.^48,66^

Electrical connections were made using a pair of silver/silver chloride electrodes (Ag/AgCl) with 1.5% agarose/2 M KCl bridges. Voltage was applied to the *cis* compartment and grounding was applied to the *trans* compartment. Thus, positive voltages indicate that the *cis*-side is positive with respect to *trans*. To minimize the impact of external noise sources, a double metal screen surrounded the bilayer chamber (Amuneal Manufacturing Corp., Philadelphia, PA). Furthermore, an anti-vibration table (Technical Manufacturing Corp. (TMC), Peabody, MA) was employed to shield the system from potential disruptions caused by mechanical vibrations. Ion channel currents were recorded at room temperature in the voltage-clamp mode using an Axopatch 200B amplifier (Molecular Devices, Sunnyvale, CA). The current was filtered at 10 kHz with an in-line low-pass 8-pole Bessel filter. The data was then digitized at a sampling rate of 50 kHz using a Digidata 1440A (Molecular Devices, Sunnyvale, CA) and transferred to a PC for analysis by pClamp 10 software (Molecular Devices, Sunnyvale, CA).

To generate current-voltage (I-V) plots, holding voltages of variable durations ranging from -120 mV to +120 mV in 10 mV steps were applied to the membrane. The amplitudes of current jumps at each applied voltage were quantified by fitting histograms of current values using a single Gaussian function.

To formally classify Tse4-currents measured with KCl in E. coli as ohmic or rectifying, we fitted the I-V plots with the following empirical equation used in Materials Science to analyze currents from semiconductor p/n diodes:^67^

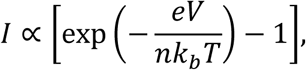

where *e* is the elementary charge, *k_B_* is the Boltzmann constant, T is the absolute temperature and *n* is an ideality factor. The lower the ideality factor (in absolute value, |*n*|), the more rectifying is the I-V curve. For instance, typical non-ohmic I-V curves from ideal semiconductor diodes yield |*n*| ∼ 1-2 and similar values are obtained when applied to rectifying protein ion channels (|*n*| ∼ 1-6).^66–69^ Thus, we categorized an I-V curve from Tse4 as rectifying when |*n*| < 10. Moreover, the sign of *n* indicates the type of rectification, with n < 0 for outward rectifying (zero or low negative currents for negative potentials, high positive currents for positive potentials) and n > 0 for inward rectifying (high negative currents for negative potentials, zero or low positive currents for positive potentials) channels.

Conductance G was obtained from the slope of each I-V plot, using the entire curve for ohmic currents and the local slope in the I = 0 region for rectifying currents.

Selectivity was evaluated by measuring the reversal potential (RP, voltage at which the current is cancelled) in experiments under a salt concentration gradient. RP was obtained either from the I-V relationship (determined by the horizontal-axis intercept) or by manually cancelling the observed current. Cation-to-anion permeability ratios were calculated from the RP values with the GHK equation^70^.

### Statistics and reproducibility

Statistical analyses were performed using GraphPad Prism 9.5 and are detailed in the figure legends.

## Data availability

The authors declare that source data supporting the findings of this study are available within the paper and its supplementary information files. **Supplementary Data 1-3** are available as supplementary files and contain the Tse5^ΔCT^-Tse4 protein sequence engineered for expression and purification of Tse4, the mass spectrometry analysis confirming the expression of Tse4, and a list of essential materials employed, respectively. Data plotted in Figures are available as supplementary material in a **Source Data file**. Uncropped and unedited gel images are included in **Supplementary Fig. 4**.

## Supplementary Information

## Acknowledgements

D.A.-J. acknowledges support from Grant PID2021-127816NB-I00 funded by MICIU/AEI/ 10.13039/501100011033 and by “ERDF/EU” and IT1745-22 / Basque Government. A.G.-M. acknowledges the financial support received from the Spanish Ministry of Universities and the Grants for the requalification of the Spanish university system for 2021-2023, financed by the European Union-Next Generation EU-Margarita Salas Modality. J.R.-P., M.Q.-M. and A.A. acknowledge support from the Spanish Government (Grant PID2022-142795NB-I00 funded by MICIU/AEI/ 10.13039/501100011033 and by “ERDF/EU”), Generalitat Valenciana (Project CIGRIS/2021/021 and Project CIAICO/2023/106) and Universitat Jaume I (Project UJI-B2022-42).

## Competing interests

The authors declare no competing interest.

## Notes

### Competing Interest Statement

The authors have declared no competing interest.

